# Metabolic remodeling of microorganisms by obligate intracellular parasites alters mutualistic community composition

**DOI:** 10.1101/2025.01.29.635536

**Authors:** Ave T. Bisesi, Ross P. Carlson, Lachlan Cotner, William R. Harcombe

## Abstract

Bacteria carry many types of obligate intracellular parasites, including plasmids and bacteriophage. During infection, these parasites redirect intracellular resources away from bacterial processes toward parasite production. Because parasite-induced metabolic changes influence host traits such as growth rate, nutrient uptake, and waste excretion, parasitic infection should alter how microbes contribute to important community and ecosystem functions. Yet there are few empirical tests of how infection shapes metabolically-mediated interactions between host and non-host species. Here, we integrated a genome-scale metabolic modeling approach with an *in vitro* obligate cross-feeding system to investigate the metabolic consequences of two intracellular parasites of *Escherichia coli*: the conjugative plasmid F128 and the filamentous phage M13. We examined the impact of these parasites on interactions between bacteria in a multispecies community composed of *E. coli*, *Salmonella enterica,* and *Methylobacterium extorquens*. Modeling predicted that parasitic infection of *E. coli* should have consequences for host growth rate and the secretion and reuptake of carbon byproducts. These theoretical results aligned broadly with *in vitro* experiments, where we found that parasitic infection changed the excretion profile of *E. coli*, inducing the net externalization of lactate. We also found that parasite-driven changes to metabolism increased the density of cross-feeding species and changes to community composition were generalizable across community interactions and some host and parasite genotypes. Our work emphasizes that microbes infected by obligate intracellular parasites have different metabolisms than uninfected cells and demonstrates that these metabolic shifts can have significant consequences for microbial community structure and function.

**IMPORTANCE:** The intracellular parasites of bacteria shape the structure and function of microbial communities in a variety of ways, including by killing their hosts or transferring genetic material. This study uses a combination of flux balance analysis and an *in vitro* system consisting of *Escherichia coli*, *Salmonella enterica*, *Methylobacterium extorquens*, and two intracellular parasites of *E. coli*, the F128 plasmid and the filamentous phage M13, to investigate how parasites change the community contributions of their hosts via metabolic remodeling. Flux balance analysis suggests that parasites change intracellular demand for different metabolic processes, leading to shifts in the identities and concentrations of compounds that infected hosts externalize into the environment. This finding is supported by experimental results, which additionally show that infection can induce the production of lactate. These findings extend our understanding of how bacteriophage and plasmids shape the structure and function of microbial communities.

## INTRODUCTION

Obligate intracellular parasites of microbes, such as bacteriophage and plasmids, are consequential drivers of microbial community composition and function [1]. The ability of bacteriophage (phage) to reduce host densities, free nutrients via lysis, and promote bacterial diversity through mechanisms such as “Kill-the-Winner” [2–4] makes them central to ecosystem processes like the viral shunt and shuttle in both aquatic [5–7] and terrestrial [8] environments. Phage and plasmids also dictate microbial community structure by facilitating horizontal gene transfer [9]. By introducing new functions or physiological niches to infected cells, horizontal gene transfer can enable new interactions between microbial species such as competitor-inhibiting toxin production or mutualistic cross-feeding [10–11]. Access to these phenotypes can have significant consequences for the coexistence and relative abundances of bacterial species [10–13].

Yet neither lysis nor horizontal gene transfer are required for parasites to shape microbial communities. Metabolic conflict between hosts and parasites is an important way that parasites can influence community structure and function. When parasites infect a host, they redirect intracellular resources away from host processes to parasite production, changing host physiology and other important traits [2, 14]. Phage infection can rewire many host metabolic pathways [2, 15–19], modulating phenotypes such as bacterial growth rate and nutrient requirements. Plasmid acquisition is often disruptive to host metabolism, as well [20]; in some cases, plasmids can change host metabolism so substantially through the induction of stress responses or cytotoxic gene products that they protect hosts against antibiotics without conferring any resistance genes [21]. Even under environmental conditions selectively favorable for traits carried by the infecting plasmid [22], metabolic carriage costs are typically sustained until the plasmid is lost during cell division [20] or some kind of compensatory evolution occurs [23]. Highly diverse parasites may even induce convergent metabolic responses, as has been shown in *Pseudomonas aeruginosa*, where infection with one of several different plasmids can each reduce bacterial excretion of citrate [24]. These changes to host metabolism are consequential to microbial communities because interactions between microbes are built on a metabolic foundation [25]. Depending on environmental conditions [26], bacteria can compete for shared nutrients [27] or cross-feed valuable metabolites [15, 28–30]. Pervasive metabolic interactions between bacteria means that parasites should have substantial community-wide consequences if infection alters key metabolic traits like host nutrient usage, waste excretion, or growth rate.

In the ocean, as many as 40% of bacterial cells are estimated to be infected by parasites at any given time [31]. Therefore, it is almost certain that infected cells with unique, parasite-induced physiologies are widespread in marine and other environments, likely contributing to community structure and function wherever bacteria are found [2]. As a result, it is important to understand how the metabolic process of chronic parasitic infection can alter metabolically-mediated interactions between host and co-occurring non-host bacterial species. Two classes of parasites in particular, conjugative plasmids and filamentous phage, can drive chronic metabolic conflict with their hosts that is sustained over both ecological and evolutionary timescales and may cascade through communities. Filamentous phage, which can be carried either episomally or as prophage [32], impose a markedly different metabolic cost than obligately lytic or lysogenic phage. This is because filamentous phage continually extrude from living cells, generally allowing for an association between a cell host and an actively replicating phage that lasts until cellular division [33–34]. These phages are consequential for several key microbial systems. For example, they have been implicated in the delayed healing of infected wounds [35], improved biofilm formation [36] and colonization of epithelial cells [37], and reduced diffusion of antibiotics [38]. Likewise, despite their metabolic burden, conjugative plasmids have been shown to provide unexpected benefits to hosts and communities [39], mediating biofilm formation and improving community resilience to perturbation in the gut [40]. Plasmid-dependent phages are also ubiquitous in nature [41–42], making the interplay between the metabolic remodeling imposed by each of these parasites an important driver of microbial communities. Both filamentous phage and conjugative plasmids change the metabolism of their hosts over sustained timescales, meaning that they are likely to have consequences for microbial composition and function across ecosystems.

To consider the impact of these two classes of parasites on microbial communities, we used a combination of a synthetic system of three interacting bacterial species - *Escherichia coli*, *Salmonella enterica*, and *Methylobacterium extorquens* (**Fig. 1A**) - and a genome-scale metabolic modeling approach focused on the nucleotide and amino acid cost of parasitic infection with the conjugative plasmid F128 and the filamentous phage M13 (**Fig. 1B**). We compared modeling-predicted changes in host metabolic processes with *in vitro* changes in community composition and function and found that infection with our parasites of interest changed the growth rate and metabolic contributions of an infected host to the microbial community in which it was embedded. Our results underscore the importance of characterizing the metabolic heterogeneity of microbes infected by intracellular parasites to successfully predict and manage complex microbial communities.

**Figure 1.**
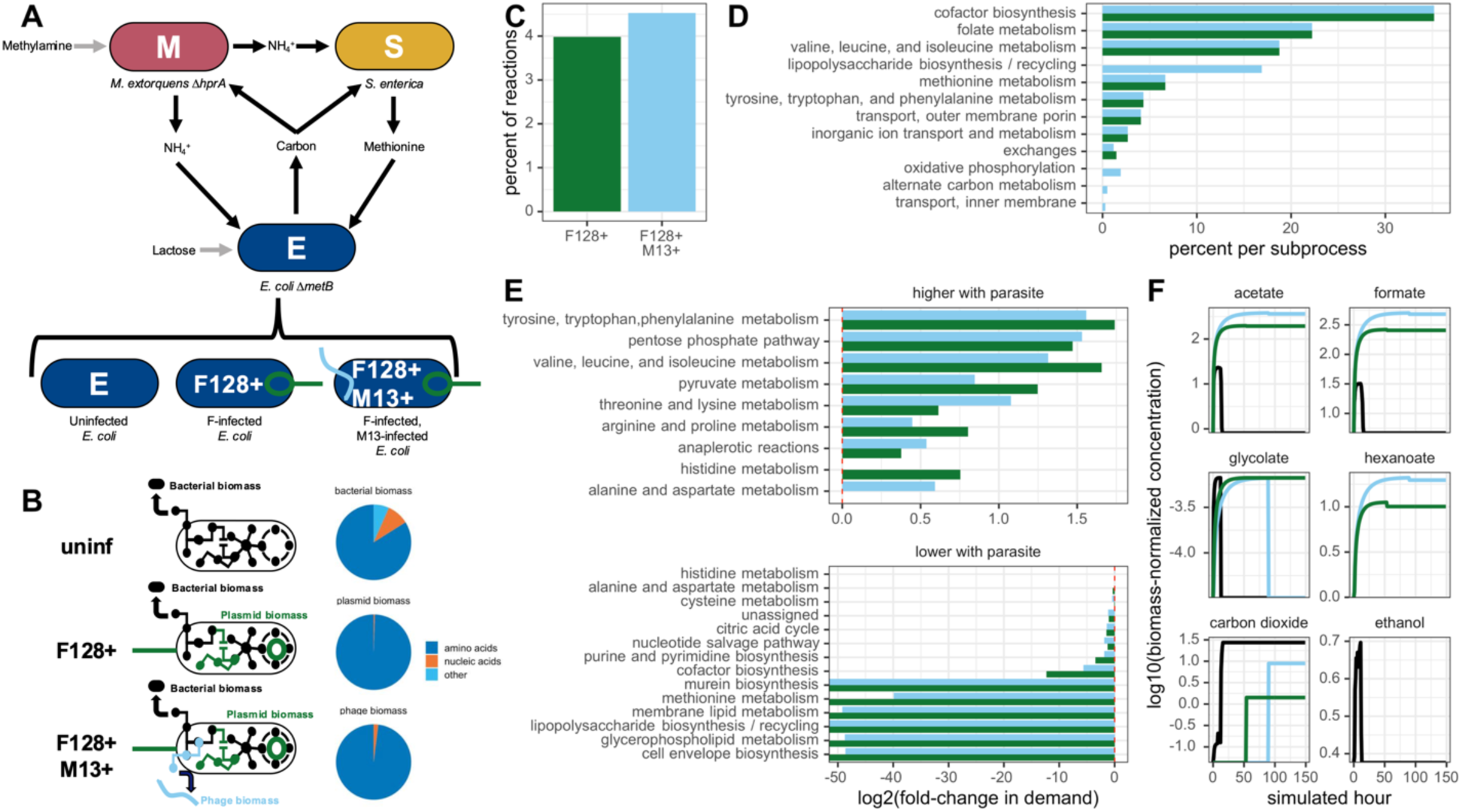
Genome-scale metabolic modeling predicts that infection increases demand for amino acids, decreases demand for host-specific processes, and limits reuptake of waste products. **A:** Schematic of synthetic microbial system with associated infection statuses of *E. coli* Δ*metB*. **B:** Schematic of *E. coli* Δ*metB* metabolic models used for FBA analyses. Models were forced to produce either only bacterial biomass or bacterial biomass in addition to plasmid and/or phage biomass. Both plasmid and phage biomass served as nutrient sinks in metabolic models; extracellular transport of phage particles was not modeled. Pie charts show the ratio of the components of biomass reactions for hosts, phage and plasmids used in FBA simulations. **C:** Percent of total metabolic reactions in the *E. coli* Δ*metB* metabolic system with non-overlapping flux ranges (FVA) when parasite biomass is optimized versus when host biomass is optimized. F128+ M13+ simulations were run with a lower bound of 0.9 mmol gDW^-1^hr^-1^ on plasmid production while maximizing M13 production. **D:** Percent of reactions in each cellular subprocess with non-overlapping flux ranges (FVA) when parasite biomass is optimized versus when host biomass is optimized. F128+ M13+ simulations were run with a lower bound of 0.9 mmol gDW^-1^hr^-1^ on plasmid production while maximizing M13 production. **E:** Log2(fold-change) in flux magnitude (i.e. demand) for all reactions in each cellular subprocess (pFBA) when parasite biomass is optimized versus when host biomass is optimized. Positive values indicate that there is more demand for a subprocess when parasite biomass is optimized. Negative values indicate that there is less demand for a subprocess when parasite biomass is optimized. F128+ M13+ simulations were run with a lower bound of 0.9 mmol gDW^-1^hr^-1^ on plasmid production while maximizing M13 production. Only cellular subprocesses where the absolute value of the log2(fold-change) was greater than 0.5 for at least one infection status are shown. **F:** Simulated log10(biomass-normalized concentrations) of key externalized metabolites during monoculture growth of *E. coli* Δ*metB* with various infection statuses (dFBA). Lactose is the only carbon source provided in the initial media. F128+ was modeled with a lower bound on plasmid production of 0.9 mmol gDW^-1^hr^-1^, and F128+ M13+ was modeled with a lower bound of 0.9 mmol gDW^-1^hr^-1^ on plasmid production and 0.06 mmol gDW^-1^hr^-1^ on phage production.

## RESULTS

### *Metabolic conflict between* E. coli *and its parasites is predicted to be driven primarily by demand for amino acids*

To predict the consequences of parasitic infection for host metabolic processes, we performed genome-scale metabolic modeling [43] to consider metabolic conflict between *E. coli* Δ*metB* and two intracellular parasites, F128 and M13 (**Fig. 1A**). Parasites were modeled using amino acid and nucleotide costs predicted by their genomic content, forcing the metabolic system of *E. coli* Δ*metB* to produce parasite biomass in addition to or instead of host biomass (**Fig. 1B**). We followed a modeling approach that has been previously applied to plasmids [44], DNA and RNA-based lytic bacteriophage [45–46], and human viruses [47]. In addition to host biomass, cells were forced to produce either F128 biomass (F128+) or both F128 and M13 biomass (F128+ M13+). Parasite biomass served as a metabolic sink for the intracellular metabolites required for production; though *in vitro* filamentous phage exit the cell continuously without lysis, we did not include extracellular transport of phage in our pseudoreactions. Additionally, we did not complete a full factorial of infection statuses because M13, like related Ff filamentous phages and many other phages, relies on the conjugative pilus to gain cellular entry to hosts [32–33, 41–42]. Infection with F128 (or any plasmid that carries the conjugative pilus machinery) is therefore a prerequisite for M13 infection in most natural systems [32–33].

With our integrated host-parasite metabolic models, we first sought to determine if differences in host metabolism arose because of infection. We did so by considering three potential metabolic scenarios using the same model: 1) a host-optimized state, where the metabolic system of *E. coli* Δ*metB* was optimized to produce only new bacterial biomass and no parasites were produced, 2) a parasite-optimized state, where the metabolic system of *E. coli* Δ*metB* was optimized to produce new parasite particles and no bacterial biomass was produced, and 3) an intermediary state, where the metabolic system of *E. coli* Δ*metB* was optimized for the production of new host biomass, but a minimum lower bound was placed on parasite production. The first two states represented two ends of a continuum of infection, while the intermediate state was more representative of the average host cell state during infection, where bacterial reproduction was possible but significantly constrained by the metabolic demands of infection. When modeling hosts infected with only the plasmid in a parasite-optimized state, a constant lower bound of 0 was placed on M13 production (lower flux bound on phage pseudoreaction = 0 mmol gDW^-1^hr^-1^), while plasmid production was optimized. When modeling hosts infected with both parasites in a parasite-optimized state, a constant lower bound was placed on plasmid production (lower flux bound on plasmid pseudoreaction = 0.9 mmol gDW^-1^hr^-1^), while M13 production was optimized. For each scenario, we investigated the full range of potential FBA solutions by performing flux variability analysis (FVA), parsimonious flux balance analysis (pFBA), and dynamic flux balance analysis (dFBA) using the existing tools CobraPy [48] and COMETS [49].

After obtaining FVA solutions, we identified metabolic sites of conflict by comparing the flux ranges of all reactions predicted by FVA between a host-optimized and parasite-optimized state. Reactions where the flux ranges did not overlap were considered high conflict, given that, by definition, they had no shared solutions between optimization states. Of the 2,585 metabolic reactions in the *E. coli* Δ*metB* metabolic system, we found that 103 (4.0%) were sites of metabolic conflict when *E. coli* Δ*metB* was infected with F128 and 117 (4.5%) were sites of metabolic conflict when *E. coli* Δ*metB* was infected with both parasites (**Fig. 1C**). All the predicted sites of conflict in F128+ except one were shared by F128+ M13+ *E. coli* Δ*metB*, while 15 reactions were unique to F128+ M13+ *E. coli* Δ*metB*, consistent with the significant amino acid and nucleotide cost of simultaneously producing plasmid and phage biomass.

We next matched each high conflict reaction to its corresponding cellular subprocess. We found that three amino acid processes - methionine metabolism, tyrosine, tryptophan and phenylalanine metabolism, and valine, leucine and isoleucine metabolism - were disrupted to a similar degree during each type of infection (**Fig. 1D**). We also saw that infection with M13 increased conflict in lipopolysaccharide, oxidative phosphorylation, and inner membrane transport reactions, consistent with the expectation that producing parasite biomass should downregulate host-specific processes (**Fig. 1D**). To understand whether these reactions were high conflict due to an increased or decreased demand for those cellular subprocesses during infection, we quantified the log2(fold-change) in parsimonious flux for all reactions in each subprocess in a parasite-optimized relative to a host-optimized state (**Fig. 1E**). Infection increased demand for four amino acid biosynthesis processes regardless of infection state, as well as the pentose phosphate pathway, pyruvate metabolism, and anaplerotic reactions (**Fig. 1E**). In contrast, infection eliminated demand for many host-specific processes, particularly cofactor biosynthesis, murein biosynthesis, and other membrane biosynthesis pathways (**Fig. 1E**). Taken together, these findings underscored that nucleotide and amino acid limitation can drive conflict between hosts and parasites and that downregulation of host-specific processes like membrane biosynthesis may occur when parasite particles do not require lipids for production.

Finally, we evaluated our intermediary state by setting lower bounds on parasite production for the F128+ model (lower flux bound on plasmid pseudoreaction = 0.9 mmol gDW^-1^hr^-1^, lower flux bound on phage pseudoreaction = 0 mmol gDW^-1^hr^-1^) and the F128+ M13+ model (lower flux bound on plasmid pseudoreaction = 0.9 mmol gDW^-1^hr^-1^, lower flux bound on phage pseudoreaction = 0.06 mmol gDW^-1^hr^-1^). These bounds were chosen because they enabled growth in all monoculture and co-culture conditions when running simulations in COMETS using dFBA (**Supplemental Fig. 1, Supplemental Table 1**). These bounds also matched the lower bound on plasmid production used for F128+ M13+ models in our pFBA and FVA analyses. With these intermediary models, we examined how differentially infected *E. coli* Δ*metB* was predicted to use and produce different compounds in monoculture. We did so by considering the biomass-normalized concentration of various media metabolites that accumulated during cellular growth due to excretion by *E. coli* Δ*metB*. We found that uninfected *E. coli* Δ*metB* produced various carbon waste products at lower concentrations per cell than infected bacteria (**Fig. 1F, Supplemental Table 1**). dFBA simulations also predicted that the reusable waste products acetate, glycolate, and formate, generated during overflow metabolism, were rapidly reassimilated by uninfected *E. coli* Δ*metB*, while F128+ and F128+ M13+ *E. coli* Δ*metB* did not take these products back up at the same rate as uninfected cells (**Fig. 1F, Supplemental Table 1**). Because acetate is an organic acid that can be used as a sole carbon source by a wide variety of microbes [50], we expected that its increased accumulation during growth of infected *E. coli* Δ*metB* could have significant implications for microbial communities. Infection also increased the per-cell production of hexanoate and altered the production of ethanol and carbon dioxide (**Fig. 1F, Supplemental Table 1**). Altogether, our genome-scale metabolic modeling results predicted that infection by F128 and M13 should alter host metabolism and drive changes in chemical exchange.

### *Parasitic infection of* E. coli *changes host growth rate and excretion profile in vitro*

Because metabolic modeling suggested that infection should change the growth rate, total yield, and effective excretion profile of *E. coli* Δ*metB* (**Fig. 1F, Supplemental Table 1**), we next sought to test these predictions using our *in vitro* system. We infected *E. coli* Δ*metB* with our parasites of interest and grew these strains in monoculture to identify potential changes in growth rate or yield (**Fig. 2A**). We found that, consistent with our dFBA modeling predictions (**Supplemental Fig. 1A**), infection reduced the growth rate of hosts (**Fig. 2B**, *P* < 0.001), though infection with both parasites did not significantly reduce growth rate relative to infection with the plasmid alone (**Fig. 2B**, *P* = 0.77). In contrast to dFBA, we observed no significant change in the productivity of infected conditions (**Fig. 2B**, *P* > 0.33, **Supplemental Fig. 1**), indicating a minimal relationship between growth rate and yield, as might be expected under a high efficiency growth strategy [51].

**Figure 2.**
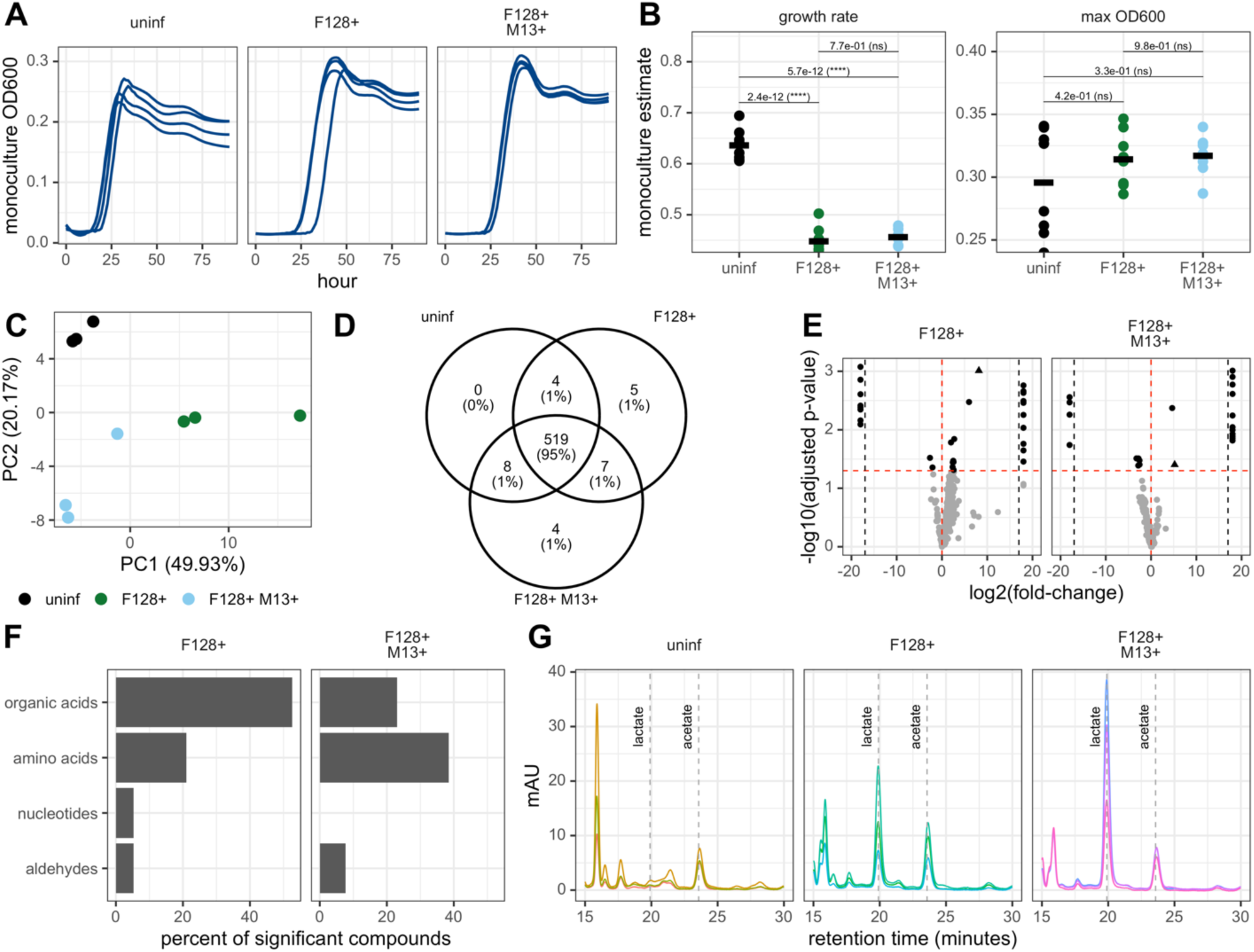
Parasitic infection of *E. coli* changes host growth rate and excretion profile. **A:** OD_600_ growth curves for *E. coli* Δ*metB* grown in monoculture. Curves are shown for a single experimental run. **B:** Estimated growth rates and maximum measured OD_600_ for *E. coli* Δ*metB* grown in monoculture. Statistical significance was determined using one-way ANOVA with Tukey’s HSD test. Data are taken from two independent experiments run during different weeks with four biological replicates per condition per experiment. **C:** PCA plot of 548 metabolic features for three spent media conditions, each with three replicates. **D:** Venn Diagram of number of metabolites detected in each of three spent media conditions across three replicates. **E:** Volcano plot of targeted metabolites from both types of infected *E. coli* Δ*metB* spent media compared to uninfected spent media. Black dots have significant log2(fold-change) values following Benjamini-Hochberg adjustment for multiple comparisons. The triangle corresponds to lactate. Points past the black dotted lines correspond with compounds with infinite log2(fold-change) values. **F:** Percent of all metabolites significantly enriched in infected *E. coli* Δ*metB* spent media compared to uninfected spent media within each compound class. Of the 9 total classes represented across enriched metabolites, only amino acid, organic acid, nucleotide and aldehyde classes are shown, as they account for the majority of enriched compounds. **G:** High-performance liquid chromatography traces for three biological replicates of spent media from *E. coli* Δ*metB* with various infection statuses. Retention times for lactate and acetate are labeled with dotted lines and text.

We next sought to understand whether infection changed the effective excretion profile of *E. coli* Δ*metB*, as dFBA predicted. We grew *E. coli* Δ*metB* with different infection statuses to mid-log and performed liquid chromatography mass spectrometry (LC-MS) using a targeted metabolomics panel on filtered spent media. Mid-log was chosen to capture the secretion of carbon byproducts prior to reuptake and optimize the level of phage infection, given that the number of phage-free cells was expected to increase as cultures approached stationary [32]. Our panel detected 548 total compounds present in at least one tested *E. coli* Δ*metB* condition. Following normalization to account for differences in biomass (as colony-forming units) at the time of sample filtration, we completed principal components analysis across all 548 features to observe the level of overlap between infection statuses (**Fig. 2C**). We found that our three infection statuses were distinguishable, though F128+ and F128+ M13+ samples had greater overlap between conditions and more replicate-to-replicate variability (**Fig. 2C**) in metabolic features than uninfected *E. coli* Δ*metB*. While infection statuses had distinct metabolomic profiles from uninfected hosts, there was still significant conservation in the identities of detected metabolites. 519 of 548 compounds were detected in all three conditions, with five compounds detected only in F128+ spent media and four compounds detected in only F128+ M13+ spent media (**Fig. 2D, Supplemental Table 2**). Seven compounds were detected in both types of infected spent media but not uninfected spent media (**Fig. 2D, Supplemental Table 2**). Given that very few compounds were detected only in phage-infected conditions, these metabolomic results were not supportive of alternative hypotheses for changes to excretion profile, such as the possibility that phage freed a wide diversity of nutrients via cellular lysis [15] or by increasing general cellular leakiness [52–56]. Instead, metabolomics suggested that differences in the excreted compounds were the result of changes in the activity of specific metabolic pathways.

To identify differences in external compounds, we calculated the log2(fold-change) in biomass-normalized concentration of all detected metabolites for infected spent media types relative to uninfected spent media. We used a t-test with Benjamini-Hochberg p-value adjustment to determine the significance of log2(fold-change) values due to multiple comparisons (**Fig. 2E**). Relatively few (<10%) of detected compounds had significantly different concentrations across conditions. Twenty-nine compounds had significantly different concentrations in F128+ spent media relative to uninfected spent media (19 compounds enriched and 10 depleted), while 22 compounds had significantly different concentrations in F128+ M13+ spent media relative to uninfected spent media (13 enriched and 9 depleted) (**Fig. 2E, Supplemental Table 3**). Seven compounds were significantly enriched in both infected conditions and no compounds were significantly depleted in both infected conditions (**Fig. 2F, Supplemental Table 3**). Most of the compounds that were enriched in infected spent media were organic or amino acids (**Fig. 2F**), two types of metabolites likely to be important for supporting the growth of other microbial species and therefore consequential for changing community composition and function. We were particularly interested in the differences in the concentration of lactic acid as it is an organic acid that can be used as a sole carbon source by the two other species in our synthetic system, *S. enterica* and *M. extorquens* [57–59].

We validated the enrichment of lactate in infected spent media by performing high-performance liquid chromatography (HPLC) on new mid-log spent media samples (**Fig. 2G**). Following normalization for biomass at the time of filter sterilization, we found that all three types of *E. coli* Δ*metB* produced comparable amounts of acetate (**Fig. 2G, Supplemental Table 4,** one-way ANOVA with Tukey’s HSD test *P* > 0.05), while lactate accumulation only occurred when *E. coli* Δ*metB* was infected and at similar concentrations regardless of infection type (**Fig. 2G, Supplemental Table 4,** t-test *P* = 0.185). Because there were no gene products carried on F128 or M13 that should have directly induced the production of lactate, the accumulation of lactate in our infected conditions instead suggested that the demands of infection shifted host metabolism to drive lactate production.

### *Parasitic infection of* E. coli *alters community composition and function*

Because we observed a reduction in growth rate and an increased accumulation of organic acids due to infection *in vitro*, we hypothesized that in our two- and three-species obligate mutualisms, infection should increase the density of partner species. We tested this hypothesis by performing co-culture experiments using our synthetic mutualism (**Fig. 1A**). We found that infection increased partner density regardless of partner identity or community complexity (**Fig. 3A**). This result was broadly consistent with dFBA predictions of species ratios in mutualistic communities given infection of *E. coli* Δ*metB* (**Supplemental Fig. 1B-C**). Additionally, in all cases except the three-species condition, *in vitro* infection with both the plasmid and phage increased partner density significantly over infection with the plasmid alone (**Fig. 3B,** *P <* 0.0001 versus *P* = 0.58). Infection also increased productivity of the whole community in some cases, generally resulting in a higher total maximum yield during growth with *M. extorquens* or both partners (**Fig. 3C,** *P <* 0.0056 versus *P* > 0.05). These results demonstrated that sustained parasitic infection of a single species was sufficient to significantly alter community composition and function via host metabolic remodeling.

**Figure 3.**
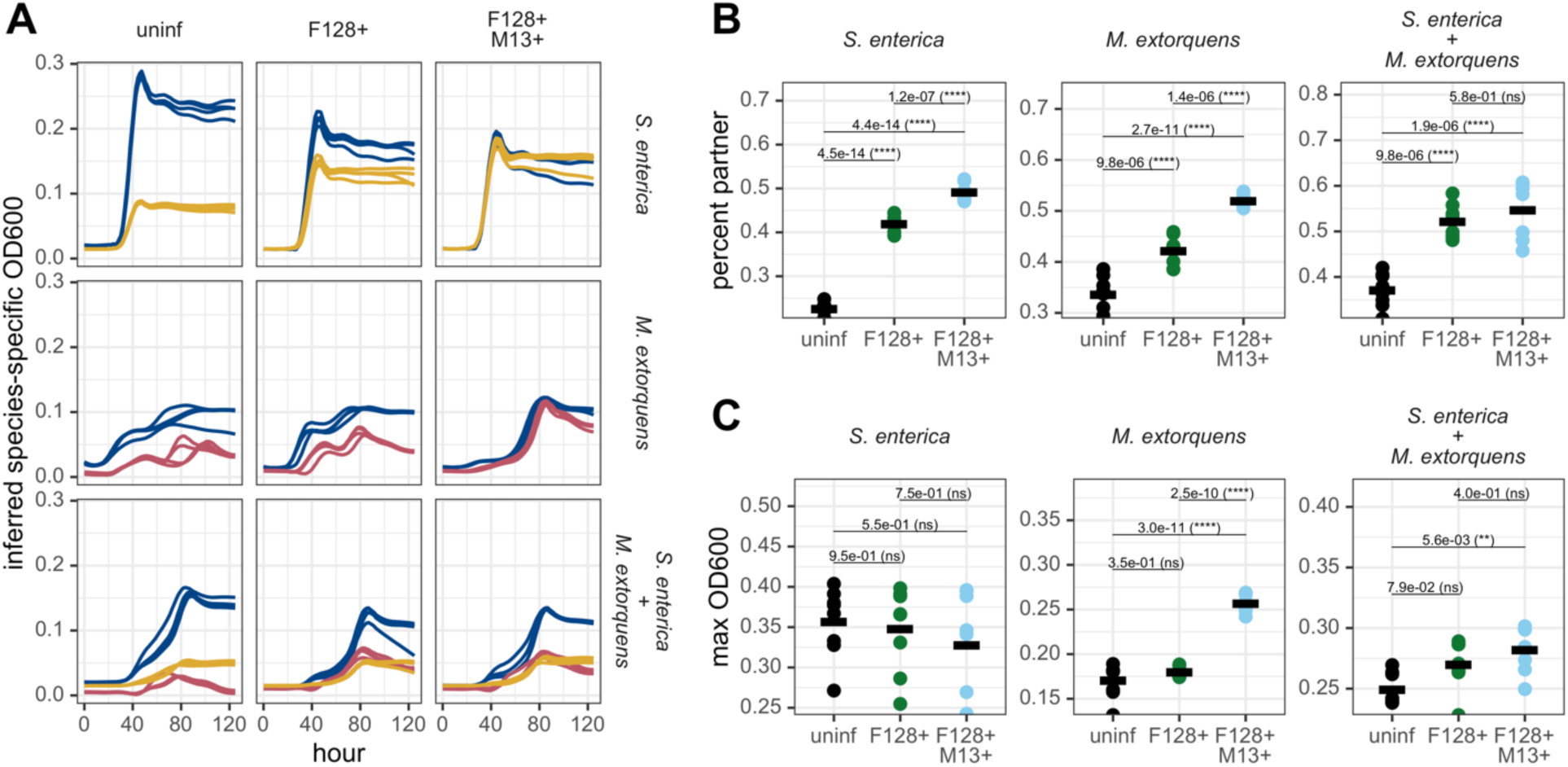
Parasitic infection of *E. coli* changes composition and productivity in co-cultures. **A:** OD_600_ growth curves for two- and three-species communities grown with *E. coli* Δ*metB* with different infection statuses. Inferred species-specific OD_600_ curves are derived from fluorescence values. Curves are shown for a single representative experimental run. **B:** Percent of co-culture composed of partner species. Statistical significance was determined using one-way ANOVA with Tukey’s HSD test. Data are taken from two independent experiments run during different weeks with four biological replicates per condition per experiment. **C:** Maximum total OD_600_ of two- and three-species communities. Statistical significance was determined using one-way ANOVA with Tukey’s HSD test. Data are taken from two independent experiments run during different weeks with four biological replicates per condition per experiment.

### *In a synthetic obligate mutualism, partner species use a combination of metabolites from* E. coli *to support growth*

Next, we attempted to determine the metabolic basis of changes in community composition. We tested what compounds were depleted by partner species from the spent media of *E. coli* Δ*metB* with different infection statuses. To do so, we grew our partner species *S. enterica* and *M. extorquens* to stationary phase in mid-log spent media from our three *E. coli* Δ*metB* infection statuses. We identified compounds depleted by partner growth by calculating the log2(fold-change) in biomass-normalized metabolite concentration and determining significance using t-tests with Benjamini-Hochberg correction for multiple comparisons. We found that only a handful of compounds were significantly depleted by our partner species (**Fig. 4A**, *P* < 0.05): 6 compounds were depleted by *S. enterica* from uninfected media, 20 compounds were depleted by *S. enterica* from F128+ media, and 7 and 6 compounds were depleted by *S. enterica* and *M. extorquens*, respectively, from F128+ M13+ media. Most of these compounds could not be used by either partner as a sole carbon source, indicating that their depletion likely supported partner growth in combination with other known system-specific compounds like acetate, galactose, and several B vitamins [60]. Glyceraldehyde - a compound present in F128+ and F128+ M13+ spent media but not uninfected spent media and an intermediate in glucose/fructose/galactose/lactose metabolism [61] - was depleted from all infected spent media conditions by both partners (**Fig. 4B, Supplemental Table 5**). Two organic acid derivatives, chlorogenic acid and 3-carboxypropyltrimethylammonium, were depleted by both partner species from F128+ M13+ spent media (**Fig. 4B, Supplemental Table 5**). Finally, aconitic acid, an intermediate in the citric acid cycle [62], was depleted by *S. enterica* from both types of infected spent media (**Fig. 4B, Supplemental Table 5**). Together, these results suggested that several differentially secreted compounds were likely to be important for supporting partner growth and that essential compounds varied between our two partner species, a finding that was broadly supported by dFBA (**Supplemental Fig. 1E-F**).

**Figure 4:**
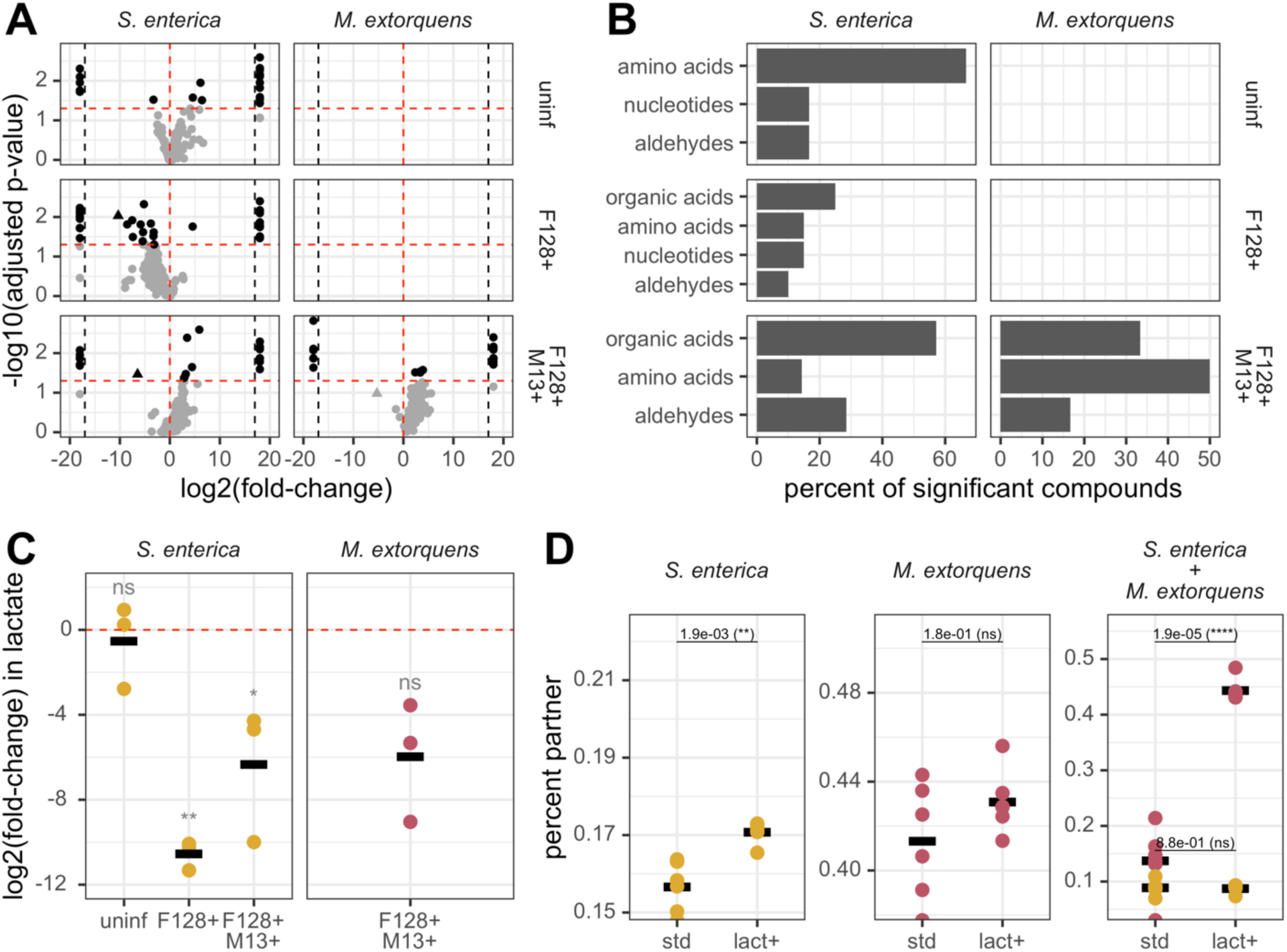
Partners use a combination of metabolites from *E. coli* Δ*metB* to support growth in mutualism. **A:** Volcano plot of targeted metabolites from both types of partner species grown in each type of *E. coli* Δ*metB* spent media. Black dots have significant log2(fold-change) values following Benjamini-Hochberg correction for multiple comparisons. The triangle corresponds to lactate. Points past the black dotted lines correspond with compounds with infinite log2(fold-change) values. No data is available for *M. extorquens* grown in uninfected or F128+ spent media. **B:** Percent of all metabolites significantly (adjusted p-value < 0.05) depleted by partner species from *E. coli* Δ*metB* spent media within each compound class. Of the 9 total classes represented across depleted metabolites, only amino acid, organic acid, nucleotide and aldehyde classes are shown, as they account for the majority of depleted compounds. No data is available for *M. extorquens* grown in uninfected or F128+ spent media. **C:** Log2(fold-change) depletion of lactic acid from all three types of *E. coli* Δ*metB* spent media by partner species. Statistical significance was determined using t-test with Benjamini-Hochberg correction for multiple comparisons across all metabolites. No data is available for *M. extorquens* grown in uninfected or F128+ spent media. **D:** Percent of final yield (maximum OD_600_) constituted by partner species during growth in obligate mutualism with uninfected *E. coli* Δ*metB* with or without 1 mmol lactate supplemented into the media. Statistical significance was determined with t-test. Data are taken from one independent experiment with six biological replicates per condition.

Given the observed accumulation of lactate, and the fact that it is one of the few compounds in our targeted metabolomics panel that could be used as a sole carbon source by both *S. enterica* and *M. extorquens* [57–59], we investigated its depletion in our metabolomics data. We found that lactate was significantly depleted by *S. enterica* from both F128+ and F128+ M13+ media (**Fig. 4C,** *P* < 0.05). There was also a large decrease in lactate from F128+ M13+ spent media following the growth of *M. extorquens*, although this change was not statistically significant following Benjamini-Hochberg correction (**Fig. 4C**, *P* = 0.105). These results bolstered our expectation that lactate production by infected *E. coli* Δ*metB* is one driver of the increased partner density and total productivity observed in our synthetic microbial community.

Lastly, to quantify the expected contribution of lactate to community composition, we grew our two- and three-species communities with uninfected *E. coli* Δ*metB* and supplemented in 1 mmol lactate, which corresponded to the expected maximum co-culture concentration of secreted lactate during infection based on our HPLC results (**Fig. 2G**). We found that the addition of lactate significantly increased *S. enterica* density in bipartite growth (**Fig. 4E**, *P* = 0.0019); however, the magnitude of this difference did not match the observed compositional differences when *E. coli* Δ*metB* was infected (**Fig. 3B**). In contrast, while lactate supplementation did not increase *M. extorquens* density in bipartite growth (**Fig. 4E,** *P* = 0.18), it did so significantly in tripartite mutualism (**Fig. 4E,** *P* < 0.0001). While there are multiple reasons that lactate addition could have different effects on community composition in comparison to increased lactate secretion [63–65], most parsimoniously, these results suggested that infection altered the exchange of a variety of metabolites in addition to lactate in our mutualistic conditions.

### *Changes in species densities during mutualistic growth with infected* E. coli *are reproducible across interaction types and host genotypes*

Finally, we investigated the generality of our experimental observation that parasitic infection can change community composition. To do so, we obtained an additional, condensed version of the conjugative F-plasmid known as pOX38 [66]. pOX38 is a 60kb plasmid that primarily encodes the conjugative machinery and a type I toxin-antitoxin system [66] (**Supplemental Table 6**), in contrast to F128, which is much larger at 230kb and carries multiple insertion sequence (IS) elements, three classes of toxin-antitoxin systems, and the lac operon in addition to the conjugative machinery (**Supplemental Table 6**). We grew *E. coli* Δ*metB* infected with pOX38 or both the plasmid and phage under conditions of obligate mutualism with *S. enterica* (**Fig. 5A**). We found that the presence of pOX38 alone did not change the density of *S. enterica* relative to growth with uninfected *E. coli* (**Fig. 5A**, *P* = 0.073), contrary to our previous finding that infection with F128 increased *S. enterica* density during co-culture growth. In contrast, infection with both pOX38 and M13 increased *S. enterica* density as previously observed in doubly infected conditions with F128 (**Fig. 5A,** *P* < 0.0001). The divergence between community composition during growth with F128+ versus pOX38+ suggested that the metabolic burden of the pOX38 plasmid was less than that of F128 (**Fig. 5A**). This finding was supported by the growth of these strains in monoculture, where infection with pOX38 significantly but modestly reduced growth rate (**Supplemental Table 7,** one-way ANOVA with Tukey’s HSD test *P* < 0.001) and did not significantly change yield relative to uninfected *E. coli* Δ*metB* (**Supplemental Table 7,** one-way ANOVA with Tukey’s HSD test *P* = 0.472). However, the use of different plasmid genotypes also underscored that M13 drove statistically significant differences in community composition regardless of co-infecting plasmid, suggesting reproducibility in the metabolic conflict imposed by the phage on the *E. coli* Δ*metB* background (**Fig. 5A-B, Supplemental Table 7**).

**Figure 5.**
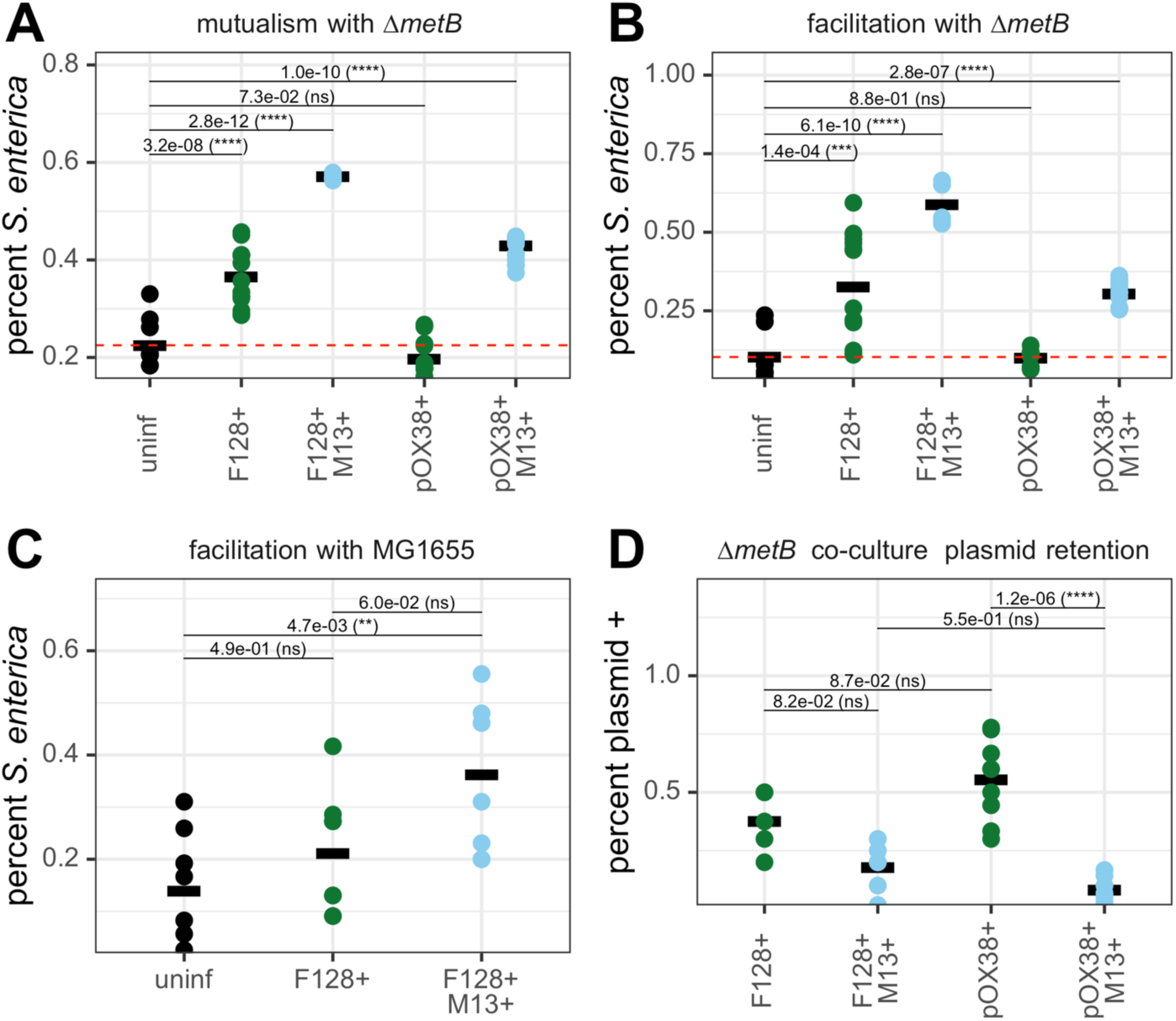
Changes in partner density occur across host genotypes, interaction types, and parasites. **A:** Percent of maximum co-culture density composed of *S. enterica* when growing in obligate mutualism with *E. coli* Δ*metB* across infection statuses. Statistical significance was determined with t-test using “uninf” as a reference group. Data are taken from three independent experiments run during different weeks with three biological replicates per condition per experiment. **B:** Percent of maximum co-culture density composed of *S. enterica* when growing in unidirectional facilitation with *E. coli* Δ*metB* across infection statuses. Statistical significance was determined with t-test using “uninf” as a reference group. Data are taken from three independent experiments run during different weeks with three biological replicates per condition per experiment. **C:** Percent of maximum co-culture density composed of *S. enterica* when growing in unidirectional facilitation with wild-type MG1655 across infection statuses. Statistical significance was determined using one-way ANOVA with Tukey’s HSD test. Data are taken from two independent experiments run during different weeks with four biological replicates per condition per experiment. Densities were calculated based on CFUs on selective media. **D:** Percent of final *E. coli* Δ*metB* population still plasmid-positive in stationary phase following growth in obligate mutualism with *S. enterica*. Densities were calculated based on CFUs on selective media. Statistical significance was determined using one-way ANOVA with Tukey’s HSD test. Data are taken from three independent experiments run during different weeks with three biological replicates per condition per experiment.

Next, we tested the impact of infection on engagement in different ecological interactions by growing *E. coli* Δ*metB* with *S. enterica* under facilitative nutrient conditions and observing the final composition of the community (**Fig. 5B**). During facilitation, *E. coli* Δ*metB* is freed from metabolic dependence on *S. enterica* due to the addition of methionine into the medium, while *S. enterica* remains dependent on *E. coli* Δ*metB* for carbon. These nutrient conditions provide a metabolically congruent environment to monoculture for *E. coli* Δ*metB*, as the strain is never under conditions of methionine limitation. We found that, consistent with our results in obligate mutualism, community composition changed significantly during infection with F128+, with increased partner density during growth with infected *E. coli* Δ*metB* (**Fig. 5B,** *P* < 0.0001). We also observed that, as in the case of obligate mutualism, infection with pOX38 alone did not change community composition relative to uninfected *E. coli* Δ*metB* (**Fig. 5B**, *P* = 0.88), while infection with both pOX38 and M13 did (**Fig. 5B**, *P* < 0.0001). Taken together, we found that phage infection increased partner density regardless of host interactions, and, while individual conjugative plasmids had reproducible effects between ecological interactions, different plasmid genotypes had different overall effects on community composition (**Fig. 5B**).

While our results suggested that changes to community composition were not generalizable across different plasmid genotypes, we also tested whether infection increased partner density regardless of host genetic background. We did so by infecting a wild-type MG1655 strain with F128 or F128 and M13. We grew these strains in lactose minimal media with *S. enterica* to replicate the nutrient conditions of facilitation with *E. coli* Δ*metB*, because MG1655 can provide carbon byproducts to *S. enterica* without depending on *S. enterica* for any nutrients. These results broadly matched our facilitative findings with *E. coli* Δ*metB* such that infection with both parasites significantly increased the density of *S. enterica* (**Fig. 5C**, *P* = 0.0047), although during facilitative growth with MG1655, *S. enterica* never dominated the population regardless of partner infection status (**Fig. 5C**). This suggested that while shifts in community composition can be qualitatively reproducible across host backgrounds, host identity likely matters for the precise metabolic manifestations of infection and the extent of compositional changes.

Finally, we tested the retention rates of our parasites during growth in obligate mutualism with *S. enterica* by quantifying the number of plasmid-positive cells at the end of growth. We found that infection with M13 increased the rates of plasmid loss relative to infection with the corresponding plasmid alone, although not significantly in the case of F128 (**Fig. 5D**, *P* < 0.001 versus *P* = 0.082). We also observed that that pOX38 appeared to be better retained than F128 when phage was not present, perhaps due to its streamlined genome and lower metabolic cost, although this effect was not significant (**Fig. 5D,** *P* = 0.087). Regardless, high rates of plasmid loss indicated that *E. coli* Δ*metB* populations were certain to be heterogeneous and, in many cases, dominated by parasite-free cells at the end of growth, suggesting that we were consistently underestimating the true metabolic burden of all three parasites and the community composition changes they could drive because we were not able to grow strains under antibiotic selection in co-culture. Taken together, our results supported previous work suggesting that diverse intracellular parasites can drive parasite- and host-specific remodeling depending on their genomic content [67], host complementarity in amino acid and nucleotide composition [2], host range [68], and lifecycle [14]. Importantly, our results also empirically demonstrated that parasites can reshape microbial communities by changing host growth rate and metabolite exchange, regardless of the precise metabolic manifestation of host remodeling.

## DISCUSSION

Our study examined how metabolic conflict between *E. coli* and two of its obligate intracellular parasites impacted microbial community composition and function in the absence of well-explored, parasite-driven mechanisms like lysis and horizontal gene transfer. We integrated a genome-scale metabolic modeling approach with *in vitro* experiments to address this question. In our modeling, we used the nucleotide and amino acid content of two parasites to impose a metabolic cost on the host cell, in alignment with previously developed methods to evaluate the metabolic consequences of infection [44–47]. Results from these models predicted that parasitic infection changed demand for amino acids, reduced host growth rate, and altered excretion profile. Our *in vitro* experiments confirmed that infection altered host growth rate and excretion profile, in particular driving the externalization of lactate. Importantly, the *in vitro* experiments clearly demonstrated that the metabolic effects of infection on one host can significantly alter the composition and function of multispecies microbial communities. Our work therefore suggests that while the genetic background of both hosts and parasites contribute to the specific manifestation of metabolic remodeling [69], the metabolic consequences of infection for host cells are sufficient enough to change host contributions to community function.

Our FBA results first predicted that infection with F128 and M13 should drive remodeling of the *E. coli* Δ*metB* metabolism. Because parasite biomass reactions were composed of nucleotides and amino acids only, we found that forcing parasite production increased amino acid biosynthesis and downregulated host-specific lipid pathways. Our modeling results aligned with existing metabolic approaches to modeling parasite infection. These models have previously been used to demonstrate that lytic phage infection reduces flux toward cell wall, lipid, and cofactor metabolism [45–46]. Additionally, modeling the RNA phage MS2 in an *E. coli* metabolic system showed that infection upregulates the pentose phosphate pathway and the biosynthesis of most amino acids while downregulating the citric acid cycle, findings which we also observed in our system despite the mechanistic and genetic differences of F128 and M13 compared to MS2 [46]. Finally, we found that our modeling approach predicted that infection of *E. coli* Δ*metB* should change host excretion profile by increasing the concentration of key organic acid byproducts like acetate produced per cell while simultaneously reducing the ability of these cells to retake up these byproducts to support additional growth.

While our modeling aligned well with existing work and predicted several key host changes *in vitro*, there are caveats to our *in silico* findings. Most obviously, FBA did not predict the observed accumulation of lactate that occurred because of infection in our *in vitro* system. We expect that our modeling predictions could be improved by imposing more biologically realistic proteomic constraints. For example, Constrained Allocation Flux Balance Analysis (CAFBA) [70] and variants of Flux Balance Analysis with Molecular Crowding (FBAwMC) [71] have been demonstrated to better reflect shifts between fermentation and respiration processes and the accumulation of resultant waste products [70–71]. Alternatively, the divergence between *in silico* and *in vitro* results could be due to a lack of metabolic optimality exhibited by infected cells (a core assumption of FBA), given that the *in vitro* infections were novel and we did not allow time for co-evolution. There is room for extensive future work to improve predictions of the metabolic impacts of novel infections.

In alignment with our modeling results, our *in vitro* study also suggested that infection with our parasites of interest reshaped *E. coli* Δ*metB* metabolism through changes to both growth rate and excretion profile. These changes had consequences for the composition and function of our two- and three-species communities regardless of partner identity or community complexity. Additionally, we found that parasite infection drove the accumulation of lactate, an organic acid with potential consequences for supporting higher densities of partner species in our synthetic microbial communities, though our results do not suggest that the induction of lactate alone was sufficient to drive the changes we observed in community composition. Nevertheless, if lactate accumulation occurs at appreciable levels in natural communities because of filamentous phage or conjugative plasmid infection and is a reproducible phenotype across hosts and parasite pairs, this could be especially important for the design and management of a wide range of microbiomes. Organic acids produced by *E. coli* during overflow metabolism have previously been shown to suppress the growth of partner species like *Pseudomonas aeruginosa* [72], and lactate, due to its acidity, can be a particularly inhibitory metabolite at high concentrations and low pH [57]. In human-associated microbiomes, lactate can also promote the growth of unwanted pathogens like *S. enterica* and *Candida albicans* [73–74]. Our results emphasize that the metabolic remodeling of bacterial hosts by obligate intracellular parasites likely serves as an important contributor to community structure and function, with consequences that may cascade beyond the microbial communities in which they are found.

However, though we observed that infected *E. coli* Δ*metB* cells produced lactate, we were not able to identify the mechanistic basis of this phenotype. There are several possibilities that deserve further investigation. For one, changes in bacterial excretion profile have been linked to changes in the ratio of cell surface area to volume (S/V) [75–76], such that an increased S/V ratio can cause increased cellular leakiness [75] and alter the dominance of respiration versus fermentation metabolic processes in *E. coli* [76]. If infection contributes to alterations in cell S/V ratio, such changes could drive differential production of lactate and other organic acids relevant to our cross-feeding system. A related possibility is that the production of the conjugative pilus and M13 extrusion channels [77–78] could push cells toward fermentative processes by reducing the enzymatic capacity of the inner membrane [72]. There is also the possibility that lactate production is related to the induction of the phage-shock protein (Psp), which, while it was originally discovered to activate in response to the M13 gene product pIV [79], can also be induced by any reduction in the energy status of the cell [80] resulting in increased inner-membrane permeability [81]. However, we expect that the most parsimonious explanation for lactate production is the fact that the proteomic cost of respiration exceeds that of fermentation, particularly in terms of nitrogen investment [82]. The metabolic burden of producing nitrogen-rich parasite particles may require metabolic strategies that increase proteomic efficiency, driving increased overflow metabolism [82]. The extent to which this phenotype is generalizable across parasite-host pairings warrants continued attention.

Additionally, our results suggested that different parasite genotypes are likely to drive different types of metabolic remodeling, although we were not able to identify the mechanistic underpinnings of the community differences between F128 and pOX38. We expect that a major contributing factor is the fact that these parasites have varying genomic content. Because conjugative plasmid-mediated IS transposition has been shown to mediate bacterial adaptation to antibiotics [83], the presence of IS elements on F128 could have promoted gene inactivation in infected *E. coli* Δ*metB* in metabolically consequential ways. Furthermore, toxin-antitoxin systems have been shown to induce metabolic stress [84], which may have contributed to more substantial remodeling of *E. coli* Δ*metB* by F128. The presence of the lac operon on F128 also represents a potential explanation. Though the F-plasmid has a narrow host range (Enterobacteriaceae) [85], infection with F128 can enable lactose utilization in *S. enterica* [86], meaning that in co-culture with F128+ *E. coli* Δ*metB*, conjugation between the two strains could have freed a subset of *S. enterica* from nutrient dependence [87–88]. Therefore, the fact that pOX38 does not carry the lac operon could explain the divergence in species ratios between these two parasite conditions. However, we did not observe the restoration of lactose utilization by *S. enterica* grown in co-culture (**Supplemental Fig. 2A**) in any of our experiments, likely because cultures were grown in liquid at small volumes. Though conjugation between *S. enterica* LT2 and our F128+ *E. coli* strain was detected at moderate rates on agar plates (**Supplemental Fig. 2B**, 5-20%), in liquid, if conjugation occurred, it did so at low rates (**Supplemental Fig. 2C**, <1%). As a result, we do not expect that transfer of the lac operon - nor the process of conjugation as a whole - was primarily responsible for the observed changes in community composition and function, though it could be a significant force in natural communities [39, 89–90]. Instead, our results emphasize that the exact metabolic manifestations of infection are likely specific to host-parasite pairs and may need to be evaluated on that basis.

To that end, our results lay the foundation for interesting future work. First, we suggest that it will be important to better understand the community-wide consequences of maintaining multiple parasites in a single host. For example, while the metabolic cost to hosts often increases with the number of plasmids carried [20, 91], in some cases positive epistasis between plasmids can reduce the cost of acquisition [92]. The extent of metabolic remodeling by a parasite and its community consequences are therefore likely to depend on the other parasites with which it co-occurs. Understanding how metabolic remodeling changes over evolutionary time is also essential to predicting and managing microbial communities. Long-term associations between parasites and hosts are often enabled by compensatory mutations that accumulate rapidly to ameliorate the costs of infection [10, 92–94]. Therefore, it is possible that coevolution between *E. coli* and the parasites tested here would minimize the extent of metabolic modeling. Because parasite infection is often transient in natural communities, we expect that our results provide a snapshot of how infection can alter species coexistence when hosts are confronted with novel parasites, though we suggest evolutionary studies would be an important next step. Finally, because poor nutrient conditions have been found to maintain filamentous phage infections by increasing the cost of resistance mutations [95], microbial community composition and parasite infection are almost certain to be engaged in feedback loops across ecosystems, where resource availability driven by cross-feeding and competition change host infection dynamics and host infection then alters nutrient concentrations by changing host metabolism.

In this study, we combined a genome-scale metabolic modeling approach with ecological experiments to explore the role of metabolic conflict between hosts and parasites in shaping community composition and function. Our work demonstrates empirically that metabolic conflict between microbial hosts and intracellular parasites is sufficient to shift the structure of the broader community and underscores the diversity of mechanisms through which parasites drive community and ecosystem function. Understanding the higher order ecological effects of host-parasite interactions is central to our ability to manage human-associated microbial communities and design microbiomes for important purposes like bioremediation. Given the ubiquity of parasite infection across all environments where microbes are found, future work should focus on identifying patterns in how parasitic infection reshapes microbial community function across host-parasite pairs, parasite-parasite interactions, and diverse ecosystems.

## MATERIALS AND METHODS

### Flux balance analysis

Flux balance analysis has been extensively described. It is a linear optimization method used to reconstruct cellular metabolic networks [43]. By assuming a steady state, FBA optimizes a set of fluxes given an objective function and bounds on the flux allowed through each reaction [43]. The FBA linear optimization problem is formalized as:

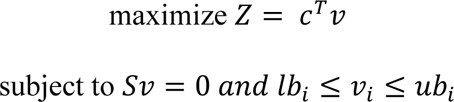

where *S* is the stoichiometric matrix and defines the stoichiometry of all metabolites across all reactions, *v* is the vector of reaction fluxes, and *c* is the vector specifying the objective function *Z* [43]. Constraints on the flux through each reaction *v_i_* are defined by imposed lower (*lb_i_*) and upper flux (*ub_i_*) bounds [43]. Using this formalization, we performed FBA, parsimonious FBA (pFBA), and flux variability analysis (FVA). For pFBA, in addition to the previously specified constraints, an additional constraint is imposed to minimize total flux through the system (i.e. minimize the sum of vector *v*). FVA provides the range of flux values for each reaction that could satisfy the linear optimization problem as it is formulated above.

All simulations were completed using variants of existing metabolic models. The host *E. coli* metabolism used in this study was based on the model iJO1366 [96]. To reproduce our in vitro system, a version of *E. coli* that is auxotrophic for methionine was created by knocking out the function of the *metB* gene in iJO1366 [97]. A methionine-secreting version of *S. enterica* was built using the STM_v1_0 metabolic network by forcing the model to generate 0.5 mmols of extracellular methionine for each gram of biomass growth and preventing reuptake by making methionine transport unidirectional [97]. Finally, an *ΔhprA M. extorquens* AM1 metabolic model was constructed as previously described [97]. Infected versions of the *E. coli* metabolic model were generated and evaluated as described below.

### Generation of parasite biomass functions

To integrate our parasites of interest into our metabolic models, we defined a pseudoreaction accounting for the stoichiometry required to produce a single parasite particle. Pseudoreactions were generated based on publicly available genome and protein sequence information for the dsDNA conjugative plasmid F128 (GenBank: NZ_CP014271) augmented with tetracycline-resistance genes from *Tn*10 (GenBank: AY528506) and the ssDNA filamentous phage M13 (GenBank: JX412914). Sequencing was performed on our *in vitro* parasite strains to confirm that their genomic and proteomic content aligned with the GenBank entries used for model integration.

In line with protocols developed for the integration of other intracellular parasites into host cell metabolic models [47], we first calculated the total moles of each nucleotide per mole of parasite particle, as appropriate for either dsDNA or ssDNA, such that:

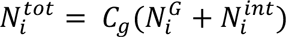

where 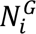 is the total number of nucleotide *i* in the parasite genome, 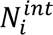 is the total number of nucleotide *i* present in any relevant replication intermediates, and their sum is multiplied by *C_g_*, the genome copy number per parasite particle, which for both parasites was a value of 1. We converted the moles of each nucleotide into grams of nucleotide per mole of parasite by multiplying 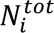 by the molar mass (g mol^-1^) of the nucleotide *i.* Finally, we expressed the stoichiometric coefficients of each nucleotide in the parasite pseudoreaction in millimoles per gram of parasite by dividing 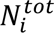 by the total molar weight of the parasite (including both nucleotides and amino acids) and multiplying that value by 1000.

The total moles of each amino acid per mole of parasite was obtained using sequence information, such that:

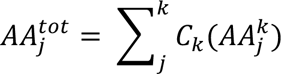

where *C_k_* is the copy number of protein *k* and 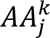 is the total number of amino acid *j* present in protein *k.* For the construction of the F128 pseudoreaction, only the 24 proteins associated with pilus formation (**Supplemental Table 7**) were assigned a copy number greater than one [98]. The copy numbers of pilus-associated proteins were calculated based on experimental findings that the pilus is composed of a 5-start helix in which there are 12.8 units of propilin protein per helical turn [99], with a typical maximum total length of 20 microns [100]. This provided a rough estimate for the maximum copy number of *traA*, the propilin component. All other pilus-associated proteins were scaled appropriately from that number based on their function in pilus-formation and conjugation (**Supplemental Table 8**) [101–108]. Amino acid investment in plasmid production could therefore be scaled with the total number of pili produced. For the purposes of our simulations, we imposed a cost of producing only a single pilus structure. Copy numbers of filamentous phage M13 proteins were obtained based on existing work detailing the structure and function of the 10 proteins encoded in the M13 genome [109–112] (**Supplemental Table 9**). We then converted the moles of amino acid per mole of parasite to grams of amino acid per mole of parasite by multiplying total moles of each amino acid per mole of parasite by the molar mass (g mol^-1^) of the appropriate amino acid. Finally, as with nucleotides, we expressed the stoichiometric coefficients of each amino acid in the parasite pseudoreaction in millimoles per gram of parasite using the total molar weight of the parasite.

We accounted for the use of ATP molecules to polymerize amino acid monomers by multiplying the expected number of ATP molecules required per peptide bond (4) [47] with the total amino acid counts, performing the calculation for each relevant protein, scaling the value by the assigned protein copy number, and summing across all proteins to generate the total number of ATP molecules required to produce all proteins associated with a parasite. Again, as before, we expressed the stoichiometric coefficients of ATP in the parasite pseudoreaction in millimoles per gram of parasite using the total molar weight of the parasite. We also accounted for the creation of PP_i_ molecules molecules formed via the nucleotide polymerization using the same process, by multiplying the expected pyrophosphate generation per nucleotide (1) [47] with the total nucleotide counts. The same calculation was applied to relevant genome intermediates, and the results were summed to generate the total number of PP_i_ molecule produced during genome polymerization for a parasite. We expressed the stoichiometric coefficients of PP_i_ in the parasite pseudoreaction in millimoles per gram of parasite using the total molar weight of the parasite. The total molar mass of the parasite was calculated using the total mass of all genome and proteome components.

Finally, we formalized the single-direction pseudoreaction based on the previous calculations of stoichiometric coefficients, such that the sum across the stoichiometry for nucleotide content, amino acid content, ATP requirements and H_2_O composed the left-hand side and the right-hand side summed across the stoichiometry of ADP, H^+^, and P_i_ generation via amino acid polymerization, plus PP_i_ generated by nucleotide polymerization. Reactions were added to our host *E. coli* metabolic model iJO1366 using COBRAPy. Initial upper and lower bounds on parasite production were set at 0 and 1000, respectively.

### Simulation parameters

We used the COBRAPy package v. 0.29.0 [48] to perform all FBA, pFBA, and FVA simulations. Generally, we assumed that carbon was the only limiting resource and therefore ran simulations under the condition that all other nutrients were present in excess (**Supplemental Table 10**). Lactose was used as the default carbon source for all experiments to replicate the conditions of our in vitro mutualistic system. To create a host-optimized state in our E. coli model, the objective function was set to the iJO1366 biomass function (BIOMASS_Ec_iJO1366_core_53p95M), with a lower bound of 0 on both F128 and M13 pseudoreactions. In an F128-optimized state, the objective function was set to the F128 biomass pseudoreaction with a lower bound of 0 on the M13 pseudoreaction. In an M13-optimized state, the objective function was set to the M13 pseudoreaction with a constant lower bound of 0.9 mmol gDW^-1^hr^-1^ on the F128 pseudoreaction. Optimization was completed with the Gurobi optimizer.

dFBA was completed using COMETS v. 0.5.2 [49] to simulate the metabolic state of our strains in mono- or co-culture. Simulations were performed in a liquid environment with an initial population size of 1e-8 gDW for each species. The objective function of each species was set to its core biomass reaction. All nutrients not provided via cross-feeding were expected to be present in excess, and the media was supplemented with 1000 mmols per base nutrient. All media conditions used are available in **Supplemental Table 10.** Simulations were performed for 200 cycles with a timestep of 1 hour. Species biomasses, media metabolite concentrations, and reaction fluxes were evaluated across the growth period (**Supplemental Fig. 1**). The lower bound on plasmid production was set at 0.9 mmol gDW^-1^hr^-1^ for both the F128+ state and F128+ M13+ state, while the lower bound on phage production in an F128+ M13+ state was set to 0.06 mmol gDW^-1^hr^-1^. Host-optimized and intermediate infection states for *E. coli* Δ*metB* were implemented as in COBRAPy. Optimization was completed with the Gurobi optimizer.

### FBA data analysis

Following completion of FVA, we identified those reactions where the range of predicted flux values from either parasite-optimized state did not overlap with the range of predicted flux values from a host-optimized state. Those reactions were then quantified as a percentage of all metabolic reactions in the system or as a fraction of the total number of reactions in the cellular subprocess to which they individually belonged. Additionally, using pFBA, we calculated the parsimonious reaction flux for each cellular subprocess in terms of a fold-change as previously described [47]. Briefly, the value can be calculated as:

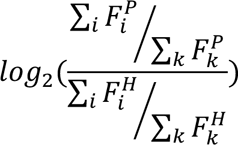

where ∑*_i_* 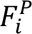 is the sum of the parsimonious flux value through all reactions that belong to cellular subprocess *i* in a parasite-optimized state, ∑*_k_* 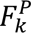 is the sum of the flux through all *k* reactions of the model in a parasite-optimized state, and the indexation *H* indicates those same values calculated when the model is in a host-optimized state. Calculations used absolute values to reflect the magnitude of changes in flux; the two reactions for which the direction of flux changed due to infection (i.e. negative to positive) were dropped from the metric. A positive log2(fold-change) value indicated higher demand for a subprocess in a parasite-optimized model, while a negative value indicated lower demand. Finally, we considered the dFBA-predicted metabolite concentrations in media from *E. coli* Δ*metB* with various infection statuses growing in monoculture. Co-culture simulations were also performed in COMETS and data are included in the supplement (**Supplemental Fig. 1**).

### Bacterial co-culture system

Our obligate mutualism has been previously described [113–114]. The bacterial strains used for these experiments are listed in **Supplemental Table 11**. Our *S. enterica* serovar Typhimurium LT2 has mutations in *metA* and *metJ* [115], causing it to oversecrete methionine. Our *E. coli* is auxotrophic for methionine due to a deletion of *metB* [113]. Lastly, our strain of *M. extorquens* AM1 can get energy from C1 compounds but cannot assimilate carbon from these sources due to a deletion of *hprA* [114]. To track individual species abundances during co-culture growth, we tagged *E. coli* Δ*metB* with a cyan fluorescent protein (CFP), *S. enterica* with a yellow fluorescent protein (YFP), and *M. extorquens* with a red fluorescent protein (RFP) [116]. A strain of MG1655 with the lac operon restored was also used in cross-feeding experiments with *S. enterica* in lactose minimal media, where MG1655 grows independently and *S. enterica* depends on carbon byproducts excreted from MG1655.

A fluorescent bacterial strain was used to confirm M13 infection. *E. coli* CSH22 (*λ−*, *thi−, trpR−,* Δ(*lac-pro*)), an F’ strain belonging to the Cold Spring Harbor Laboratory collection, was transformed with the plasmid pJS001, which induces red fluorescence when M13 particles enter the cell [94]. This strain is referred to as JS002. Because CSH22 is a proline auxotroph, the F’ plasmid is required for growth in media lacking proline, while pJS001 carries an ampicillin resistance gene. Both plasmids can therefore be maintained in JS002 during growth in minimal media without proline and with added ampicillin. Additional strain and plasmid information is available in **Supplemental Table 10**. Both strains were acquired from J. Shapiro.

### Parasite strains

The two primary parasites used were the conjugative F128 plasmid and the filamentous phage M13. The plasmid F128 (*λ−, thr-1, araC14, leuB6*(Am), *lacY1*, *glnX44*(AS), *galK2*(Oc), *galT22*, Δ*trpE63, xylA5*, *mtl-1*, *thiE1*) is a 230kb fertility factor plasmid that was acquired from the Coli Genetic Stock Center at Yale University via J. Shapiro [117]. The plasmid has a Tn*10* integration that allows the use of tetracycline to select for plasmid-positive cells. J. Shapiro provided a phage stock of M13. Sequence information for F128 and M13 can be found in **Supplemental Tables 12** and **13**, respectively. Follow-up experiments were completed with a 60kb version of the conjugative F plasmid consisting only of the conjugative machinery and the tetracycline resistance cassette called pOX38 [66]. This plasmid was provided by P. Christie; sequence information is available in **Supplemental Table 14** [66]. Unique gene products in each plasmid strain can be found in **Supplemental Table 6**.

Strains infected with F128 were obtained via conjugation with an MG1655 F128 donor. Colonies infected with both F128 and M13 were obtained by spotting 6 10µL spots of purified M13 stock onto a top agar assay of exponentially-growing cells that were already infected by the plasmid. Cells from one turbid clearing were scraped off the plate, resuspended in saline, and streaked onto an LB agar plate containing x-gal (5-bromo-4-chloro-3-indolyl-β-D-galactopyranoside) and 10µg/mL of tetracycline. Infection of individual colonies was confirmed prior to use in experiments as described below. The same procedures were performed to obtain cells infected with pOX38 and pOX38 and M13. The original pOX38 conjugative donor was a MC4100 *E. coli* strain provided by P. Christie [66].

### Media conditions

Minimal hypho liquid media was prepared as previously described [118]. Each media component was sterilized prior to mixing at room temperature [118] (**Supplemental Table 15** and **16**). As needed, solutions containing sulfur, nitrogen, phosphorus, calcium chloride, and metals were supplemented into each media type (**Supplemental Table 15** and **16**), in addition to the appropriate carbon source and methionine/methylamine. In *E. coli* Δ*metB* monoculture conditions and in ES cross-feeding media, the carbon source served as the limiting nutrient. In EM and ESM cross-feeding media, the amount of total nitrogen available in the medium (as methylamine, then metabolized by *M. extorquens*) limited total co-culture density. As a result, the final possible yield across different media types varied. These constraints were reflected in the media used for our dFBA simulations, as well. We performed routine culturing and plating of all bacterial strains with Miller Lysogeny Broth (LB) unless otherwise indicated. When doing routine culturing of any strains infected with F128 or pOX38 (including those with phage), 10µg/mL of tetracycline was added to prevent plasmid loss. When culturing strains with pJS001 plasmid, 100µg/mL of ampicillin was added to prevent plasmid loss.

### Phage-screening assays

Phage infection confirmation assays were performed in 96-well flat bottom plates on a Tecan Infinite Pro200 plate reader for 24 hours at 37°C with shaking at 432 rotations per minute. Individual phage-infected colonies streaked from plaques were resuspended from an agar plate into 200µL of saline. 5µL of each resuspended colony was then seeded into three wells containing 200µL of LB and tetracycline media, in addition to two wells containing 200µL of glucose and methionine media. 5µL of a JS002 colony suspended in 200µL of saline was also added to glucose and methionine wells along with phage-infected cells or into wells containing 200µL of glucose and ampicillin media. Up to ten infected colonies could be screened for infection at a time. After 24 hours of incubation, during which OD_600_ and red fluorescence measurements (Ex: 544, Em: 590) were taken every 20 minutes, JS002-only wells were used to establish an RFP baseline. Any wells containing both phage-infected cells and JS002 in which the RFP was at least greater than 1.5x times the RFP baseline were considered truly M13-positive. The colony with the highest relative RFP was chosen for additional experiments. LB wells seeded with cells from the chosen infected colony were then transferred into a flask containing 5mL of LB and tetracycline and incubated for at least 30 minutes at 37°C with shaking at 120rpm prior to addition in experiments.

### Community composition experiments

Community composition experiments were performed in 96-well flat bottom plates on a Tecan Infinite Pro200 plate reader for at least 96 hours at 30°C with shaking at 432 rotations per minute. Experiments were terminated when all conditions reached stationary phase. Overnight stationary phase cultures in LB (*S. enterica, E. coli* Δ*metB*) or succinate and methylamine minimal media (*M. extorquens*) started from single colonies were washed three times in saline and adjusted to a density of 10^7^ cells per mL. Relevant M13-infected colonies were screened for community composition experiments as previously described. Exponentially-growing cultures of plasmid and phage-infected *E. coli* in LB and tetracycline were grown to mid-log (∼0.5 OD_600_), washed three times in saline, and adjusted to a density of 10^7^ cells per mL. Cells were used to inoculate 200µL of appropriate medium with 3.0 x 10^5^ total cells per well (i.e. 3.0 x 10^5^ total *S. enterica* cells in monoculture; 1.0 x 10^5^ total *S. enterica* cells, 1.0 x 10^5^ total *E. coli*, and 1.0 x 10^5^ total *M. extorquens* cells in tripartite co-cultures). Starting cell densities were confirmed using colony-forming units at the start of the experiment on appropriate selective media.

We measured and recorded OD_600_, *E. coli* Δ*metB*-specific CFP (Ex: 430 nm; Em: 480 nm), *S. enterica*-specific YFP (Ex: 500 nm; Em: 530 nm), and *M. extorquens*-specific RFP fluorescence (Ex: 560 nm, Em: 610 nm) every 20 minutes. As previously described, we converted fluorescent protein signals to species-specific OD_600_ equivalents [119]. Full factorial experiments testing all relevant monocultures and co-culture conditions were completed with three to five biological replicates per condition. To protect against edge effects, only water was placed in the outer ring of the 96-well plate. A minimum of two experiments per experiment type were set up and completed during different weeks to confirm the repeatability of the results and protect against batch effects.

In experiments where one or multiple strains did not possess a fluorescent reporter, OD_600_ measurements were taken during growth and relative densities were assessed using colony-forming units plated on appropriate selective media at the termination of experiments. Rates of plasmid loss were assessed at the end of experiments by plating infected *E. coli* from both mono- and co-culture conditions onto tetracycline plates in addition to antibiotic-free plates.

### Spent media preparation and metabolomic analysis

To generate spent media for metabolomic analysis, uninfected *E. coli* Δ*metB*, *E. coli* Δ*metB* infected with F128, and *E. coli* Δ*metB* infected with F128 and M13 were prepared in exponential cultures and washed three times in saline as previously described. Cells were seeded at an OD_600_ of 0.0005 in 50mL of lactose and methionine hypho media and allowed to grow at 30°C shaking at 120rpm until mid-log (∼0.3-0.7 OD_600_). Cultures were spun down and then filter-sterilized. Final CFUs of spent media cultures were assayed on appropriate selective media prior to spinning to determine cell density during mid-log growth. Three biological replicates were generated for each infection status. 5mL of spent media from each replicate for each condition was used to grow either *S. enterica* or *M. extorquens* (for *M. extorquens,* only growth in spent media from *E. coli* Δ*metB* infected with F128 and M13 was completed). Partner strains were grown to stationary phase and cell culture media was filter-sterilized. 1mL of each condition (mid-log *E. coli* Δ*metB* spent media or stationary phase partner media) was pipetted into 1.5mL microcentrifuge tubes and placed on dry ice. Samples were sent to Metware Biotechnology Inc. for liquid chromatography mass spectrometry. A proprietary metabolomic panel of relevant metabolites was used for metabolite identification. Across all samples, a total of 589 metabolites were detected in at least one condition (**Supplemental Table 17**).

Metabolomic data were analyzed in two main ways: principal components analysis and calculations of differential presence across conditions. Prior to PCA computation, metabolite concentrations for each replicate of the three *E. coli* Δ*metB* infection conditions were normalized by the number of colony-forming units detected at the time of filter-sterilization. Data were then pre-processed using signal drift and batch effect correction, probabilistic quotient normalization, missing values imputation using k-nearest neighbors and variance stabilization through generalized logarithm (glog) transformation. Metabolic features that were not detected in any of the three *E. coli* Δ*metB* conditions were removed prior to PCA computation. These steps were performed using the pmp v. 1.6.0 R package. Calculations of differential presence were completed using the biomass-normalized concentrations for each of three replicates for each condition. Data were subsetted as appropriate based on whether calculations were examining differential concentration between infection statuses of *E. coli* Δ*metB* or differential concentration between an *E. coli* Δ*metB* condition and its corresponding partner-depleted condition. Statistical significance was determined using t-tests with Benjamini-Hochberg adjustment for multiple comparisons.

Spent media was prepared for high-performance liquid chromatography (HPLC) assays as previously described in 5mL volumes. For HPLC, samples generated from uninfected *E. coli* Δ*metB*, *E. coli* Δ*metB* infected with F128, and *E. coli* Δ*metB* infected with F128 and M13 were run on a Column Aminex HPX-87H at 46°C, flow 0.4 mL/min, solvent 1.5 mmol H_2_SO_4_, detector SPD-10A at 210 nm. Spent media data were analyzed using the R package chromatographR v. 0.7.0 for data pre-processing, alignment, and peak detection and integration. Concentrations of acetate and lactate in filtered spent media were estimated based on the linear relationship between peak area and mmol concentration using standards ranging from 0.39 mmol to 100 mmol. All spent media samples were stored at 4°C in light-protective boxes when not in use. Samples were analyzed by metabolomics or HPLC within two weeks of collection.

### Genome isolation and sequencing

M13 stocks were prepared for sequencing following ssDNA isolation. 200µL of a high titer phage lysate was combined in a 1.5mL microcentrifuge tube with 20µL of DNase Buffer I 10x (Invitrogen), 2µL DNase I (Invitrogen) and 0.4µL RNase A (Qiagen). The tube was incubated at 37°C for 1.5 hours without shaking. 8µL of 0.5M EDTA was then added to the tube, and DNase I was heat inactivated at 75°C for 10 minutes, before the addition of 0.5µL of Proteinase K and incubation at 56°C for 1.5 hours without shaking. DNA was then purified using the DNeasy Blood and Tissue Kit (Qiagen), quantified on a Nanodrop, and sent to Plasmidsaurus (https://plasmidsaurus.com/) for standard whole plasmid sequencing. Our laboratory M13 strain was found to be genetically identical to the “Rutgers” strain (GenBank: JX412914; **Supplemental Table 13**).

The F128 and pOX38 plasmids were prepared for sequencing following dsDNA isolation from plasmid-infected *E. coli* Δ*metB*. DNA was extracted from pelleted cells from an overnight culture using the DNeasy Blood and Tissue Kit (Qiagen), quantified on a Nanodrop, and sent to Plasmidsaurus (https://plasmidsaurus.com/) for standard bacterial genome sequencing. F128 was found to be identical to the DHB4 plasmid F128-(DHB4) (GenBank: NZ_CP014271), with the addition of tetracycline resistance genes from *Tn*10 (GenBank: AY528506) (**Supplemental Table 12**). pOX38 was found to match the MG1655 strain plasmid pOX38 (GenBank: NZ_OQ683454.1) (**Supplemental Table 14**).

## Supporting information

Supplemental Material Descriptions

## ACKNOWLEDGMENTS

The authors thank J. Shapiro and P. Christie for providing phage, plasmid and bacterial strains. The authors would also like to thank X. Xiong, J. N. V. Martinson, H. Ahmed, and A. Lee for helpful comments on the manuscript and experimental design. This work was supported by NSF 2226051 and NSF 2019304 to WRH. Targeted metabolomics was performed by Metware Biotechnology Inc.

## DATA AVAILABILITY STATEMENT

Data analysis, statistics, and figure generation were performed using R v. 4.2.1 using custom scripts available at https://github.com/bisesi/Phage-Community-Metabolism. Scripts for genome-scale metabolic modeling performed in Python 3.9.12 using CobraPy and COMETS, along with raw experimental data for all detailed experiments, are available at the same link.

