## Supplemental Figures 1-2 for "Metabolic remodeling of microorganisms by obligate intracellular parasites alters mutualistic community composition"

SUPPLEMENTAL FILE 2

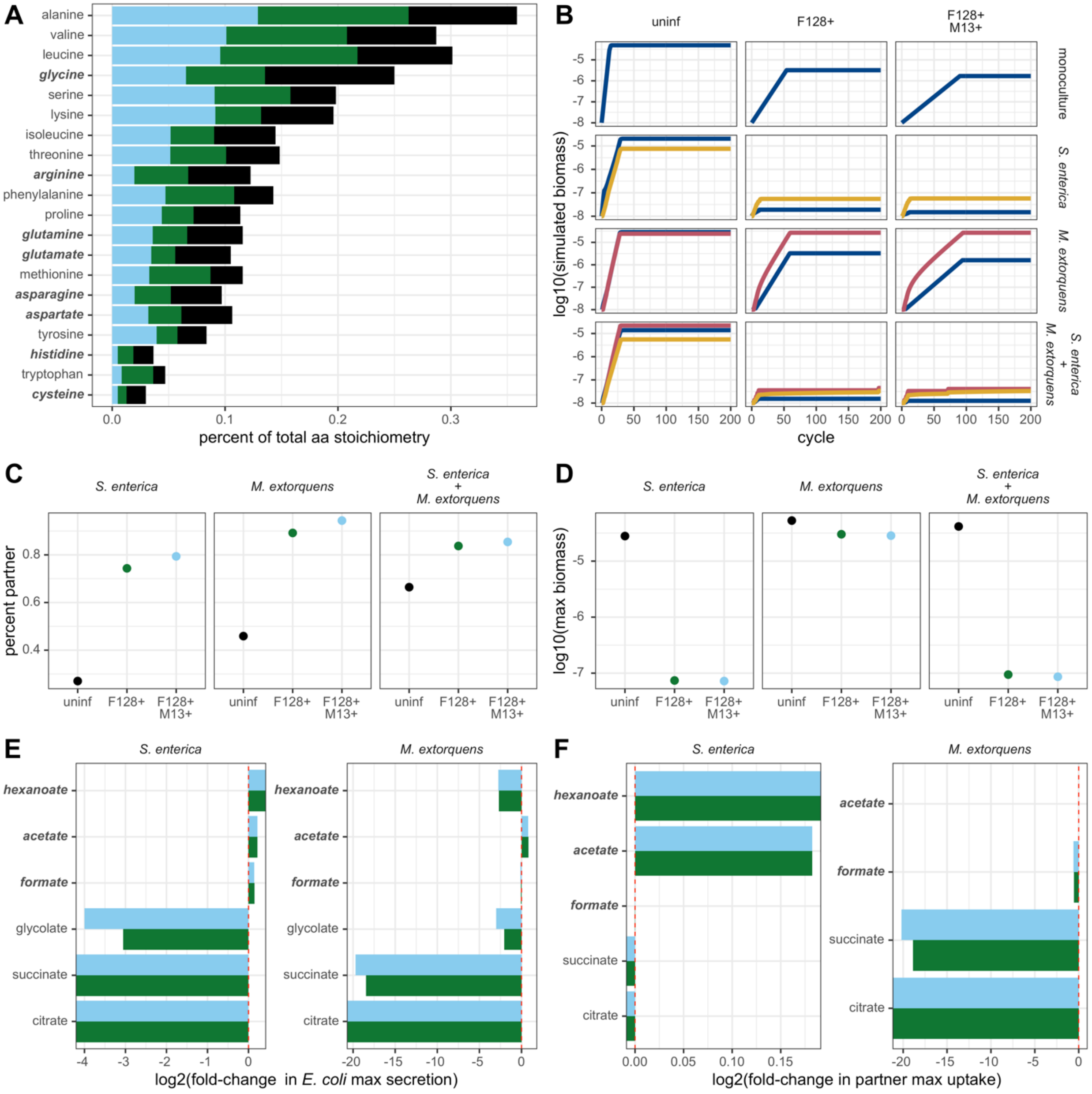


**Supplemental Figure 1. Parasitic infection of *E. coli* changes dFBA predictions of composition and productivity in co-cultures. A:** Percent of total amino acid stoichiometry represented by each amino acid for host, plasmid and phage biomass reactions. Percents are calculated by dividing the stochiometric coefficient of a single amino acid by the sum of the stochiometric coefficients of all 20 amino acids for a given organism. The names of amino acids that are a greater percentage of host stoichiometry than either parasite stoichiometry are bolded and italicized on the y-axis. **B:** Log10-transformed dFBA-simulated $OD_{600}$ growth curves for monocultures and two- and three-species communities grown with *E. coli* Δ*metB* with different infection statuses. **C:** Percent of co-culture composed of partner species at the end of dFBA co-culture simulations. **D:** Log10(maximum total simulated $OD_{600}$) for co-cultures at the end of dFBA co-culture simulations. **E:** Log2(fold-change) in maximum secretion (exchange flux) of various metabolites by *E. coli* Δ*metB* during bipartite mutualistic growth with either *S. enterica* or *M. extorquens* in dFBA simulations. Compounds that are secreted more by *E. coli* Δ*metB* during infection in at least one condition and are also taken up by either partner at a greater rate when growing with at least one type of infected *E. coli* Δ*metB* are bolded and italicized on the y-axis. **F:** Log2(fold-change) in maximum uptake (exchange flux) of various metabolites by *S. enterica* or *M. extorquens* during bipartite mutualistic growth with *E. coli* Δ*metB* in COMETS simulations. Compounds that are secreted more by *E. coli* Δ*metB* during infection in at least one condition and are also taken up by either partner at a greater rate when growing with at least one type of infected *E. coli* Δ*metB* are bolded and italicized on the y-axis. *For all dFBA simulations, F128+ is modeled with a lower bound on plasmid production of 0.9 mmol* $gDW^{-1}hr^{-1}$*, and F128+ M13+ is modeled with a lower bound on plasmid production of 0.9 mmol* $gDW^{-1}hr^{-1}$ *and a lower bound on phage production of 0.06 mmol* $gDW^{-1}hr^{-1}$*.*

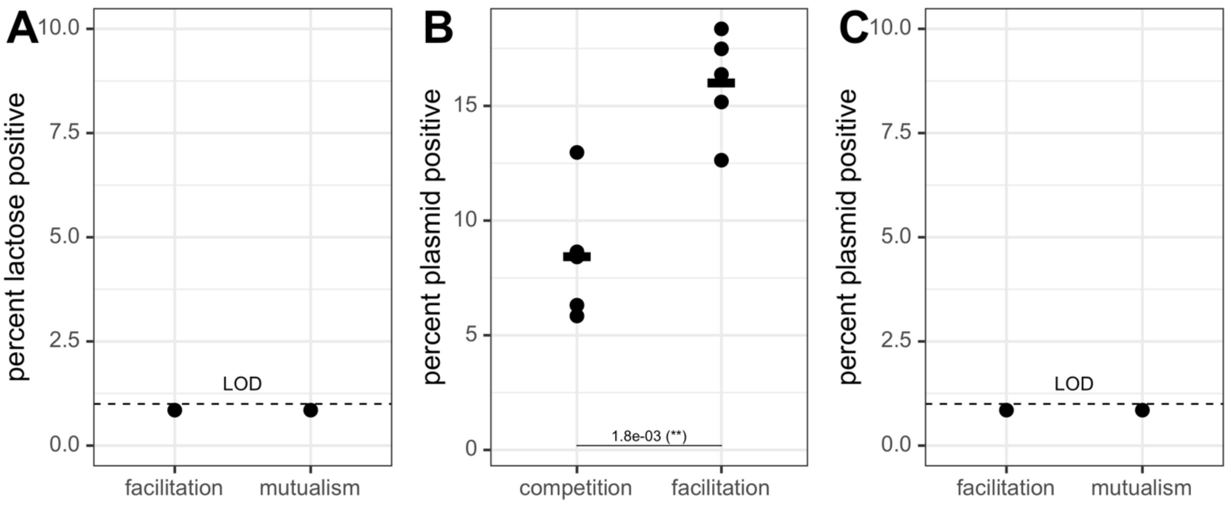


**Supplemental Figure 2. Conjugation of the F128 plasmid between *E. coli* Δ*metB* and *S. enterica* LT2 is possible on solid media but was not detected in mutualistic or facilitative experiments in liquid. A:** Lactose-positive *S. enterica* was not detected at the end of any co-culture experiments following growth with F128+ *E. coli* Δ*metB*. To test for *S. enterica* lactose utilization, half of the final co-culture volume following mutualistic growth between *S. enterica* and F128+ *E. coli* Δ*metB* (100µL) was serially diluted and plated on both lactose minimal media plates with x-gal and lactose minimal media plates with x-gal and 10µg/mL of tetracycline. Only *S. enterica* with growth enabled by infection with F128 would be able to grow on these plates. The limit of detection for plasmid-positive cells was 1% of final *S. enterica* CFU/mL. **B:** Rates of conjugation between F128+ *E. coli* Δ*metB* and a wild-type *S. enterica* LT2 on agar plates varied depending on the nutrients provided. Facilitative growth on lactose and methionine media led to a higher rate of conjugation than competitive growth on glucose and methionine media. Conjugation experiments were performed by squirting 10µL of *E. coli* Δ*metB* F128+ directly on top of 10µL of a wild-type *S. enterica* LT2 onto the surface of plates. Conjugation plates were incubated for 48 hours at 37°C before cells were scraped off plates into saline, pelleted, washed, serially diluted and plated at a volume of 100µL on either citrate minimal media plates or citrate minimal media plates with 10µg/mL of tetracycline. All *S. enterica* LT2 (but not *E. coli* Δ*metB* F128+) could grow on citrate minimal media plates, while only *S. enterica* LT2 infected with F128 could grow on citrate and tetracycline plates. The number of colonies counted on these plates were used to calculate rates of conjugation. Statistical significance was determined using a t-test. Five replicates were completed per condition. **C:** There was no detected conjugation between *E. coli* Δ*metB* F128+ and a wild-type *S. enterica* LT2 in liquid media. Conjugation experiments were performed by adding 100µL of *E. coli* Δ*metB* F128+ to 100µL of a wild-type *S. enterica* LT2 in 5mL of liquid lactose minimal media or lactose and methionine minimal media. Conjugation liquid cultures were incubated for 48 hours at 37°C before cells were pelleted, washed, serially diluted, and plated at a volume of 100µL on either citrate minimal media plates or citrate minimal media plates with 10µg/mL of tetracycline. All *S. enterica* LT2 (but not *E. coli* Δ*metB* F128+) could grow on citrate minimal media plates, while only *S. enterica* LT2 infected with F128 could grow on citrate and tetracycline plates. The limit of detection for plasmid-positive cells was 1% of final *S. enterica* LT2 CFU/mL. Three replicates were completed per condition.
