## Supplemental Material Descriptions for "Metabolic remodeling of microorganisms by obligate intracellular parasites alters mutualistic community composition"

**Supplemental File 1.** Xlsx file containing **Supplemental Tables 1-17.**

- **Supplemental Table 1:** Biomass, growth rate and maximum mmol concentration of media components secreted by *E. coli* Δ*metB* in monoculture COMETS (dFBA) simulations.
- **Supplemental Table 2:** List of metabolomic compounds not detected in uninfected spent media.
- **Supplemental Table 3:** List of metabolomic compounds significantly enriched or depleted in infected spent media relative to uninfected spent media.
- **Supplemental Table 4:** Retention times, estimated peak area, calculated concentrations and biomass-normalized acetate and lactate production by *E. coli* Δ*metB* measured by HPLC.
- **Supplemental Table 5:** List of metabolomic compounds significantly depleted by partner species from each type of *E. coli* spent media.
- **Supplemental Table 6:** Gene products unique to either the F128 or pOX38 version of the F-plasmid.
- **Supplemental Table 7:** Biomass yield (as maximum OD600) and growth rate of uninfected, pOX38+ and pOX38+ M13+ *E. coli* Δ*metB* during growth in monoculture.
- **Supplemental Table 8:** Amino acid cost of pilus production in genome-scale metabolic models. The amino acid cost of the rest of the plasmid, along with its nucleotide cost, were calculated from GenBank reference NZ_CP014271 with a gene copy number of one as described in the *Methods*.
- **Supplemental Table 9:** Amino acid cost of phage infection in genome-scale metabolic models. The amino acid cost was calculated from GenBank reference JX412914 as described in the *Methods*.
- **Supplemental Table 10:** Media conditions for dFBA (COMETS) simulations.
- **Supplemental Table 11:** Parasite and bacterial strains used for all *in vitro* experiments.
- **Supplemental Table 12:** Full list of genes and gene products in F128, NCBI accession number NZ_CP014271. The version of the F128 plasmid used in this study also encodes tet(B) and tetR(B).
- **Supplemental Table 13:** Full list of genes and gene products in M13, NCBI accession number JX412914.
- **Supplemental Table 14:** Full list of genes and gene products in pOX38, NCBI accession number NZ_OQ683454.
- **Supplemental Table 15:** Hypho minimal media conditions for monocultures.
- **Supplemental Table 16:** Hypho minimal media conditions for mutualistic and facilitative communities.
- **Supplemental Table 17:** Complete list of compounds detected in at least one sample by targeted metabolomics.

**Supplemental File 2.** PDF file containing **Supplemental Figures 1-2** with legends and captions**,** including associated methodological details.
